# Scalable Bias-corrected Linkage Disequilibrium Estimation Under Genotype Uncertainty

**DOI:** 10.1101/2021.02.08.430270

**Authors:** David Gerard

**Affiliations:** Department of Mathematics and Statistics, American University, Washington, DC, 20016, USA

**Keywords:** attenuation bias, genotype likelihood, linkage disequilibrium, polyploidy, reliability ratio

## Abstract

Linkage disequilibrium (LD) estimates are often calculated genome-wide for use in many tasks, such as SNP pruning and LD decay estimation. However, in the presence of genotype uncertainty, naive approaches to calculating LD have extreme attenuation biases, incorrectly suggesting that SNPs are less dependent than in reality. These biases are particularly strong in polyploid organisms, which often exhibit greater levels of genotype uncertainty than diploids. A principled approach using maximum likelihood estimation with genotype likelihoods can reduce this bias, but is prohibitively slow for genome-wide applications. Here, we present scalable moment-based adjustments to LD estimates based on the marginal posterior distributions of the genotypes. We demonstrate, on both simulated and real data, that these moment-based estimators are as accurate as maximum likelihood estimators, and are almost as fast as naive approaches based only on posterior mean genotypes. This opens up bias-corrected LD estimation to genome-wide applications. Additionally, we provide standard errors for these moment-based estimators. All methods are implemented in the `ldsep` package on the Comprehensive R Archive Network https://cran.r-project.org/package=ldsep.

## 1 Introduction

Pairwise linkage disequilibrium (LD), the statistical association between alleles at two different loci, has applications in genotype imputation [Wen and Stephens, 2010], genome-wide association studies [Zhu and Stephens, 2018], genomic prediction [Wientjes et al., 2013], population genetics [Slatkin, 2008], and many other tasks [Sved and Hill, 2018]. LD is often estimated from next-generation sequencing technologies, where the genotypes and haplotypes are not known with certainty [Gerard et al., 2018]. Thus, researchers typically use estimated genotypes, such as posterior mean genotypes [Fox et al., 2019], to estimate LD. However, this can cause biased LD estimates, attenuated toward zero, implying loci are less dependent than in reality. This bias is particularly strong in polyploids, and so in Gerard [2021] we derived maximum likelihood estimates (MLEs) that have lower bias and are consistent estimates of LD.

Unfortunately, the MLE approach is prohibitively slow. Researchers typically calculate pairwise LD at genome-wide scales, and the MLE approach takes on the order of a tenth of a second. Thus, for many genome-wide applications, containing millions of SNPs, LD estimation using the MLE approach would take years of computation time. This is not conducive to large-scale applications.

Here, we derive scalable approaches to estimate LD that account for genotype uncertainty (Section 2). Our methods use only the first two moments of the marginal posterior genotype distribution for each individual at each locus, which are often provided or easily obtainable from many genotyping programs. We calculate sample moments from these posterior moments, and use these to multiplicatively inflate naive LD estimates. We show, through simulations (Section 3.1) and real data (Section 3.2), that our estimates can reduce attenuation bias and improve LD estimates when genotypes are uncertain. All calculations have computational complexities that are linear in the sample size, and so these estimates are scalable to genome-wide applications.

## 2 Methods

In this section, we will define moment-based estimators of the LD coefficient ∆ [Lewontin and Kojima, 1960], the standardized LD coefficient ∆′ [Lewontin, 1964], and the Pearson correlation *ρ* [Hill and Robertson, 1968]. We will only consider estimating the “composite” versions of these LD measures which, advantageously, are appropriate LD measures for generic autopolyploid, allopolyploid, and segmental allopolyploid populations, even in the absence of Hardy-Weinberg equilibrium [Gerard, 2021]. We will also only consider biallelic loci, where the genotype for each individual is the dosage (from 0 to the ploidy) of one of the two alleles.

We wanted to create LD estimators that account for genotype uncertainty while also being agnostic to the genotyping technology (e.g., microarrays [Fan et al., 2003], next-generation sequencing [Baird et al., 2008, Elshire et al., 2011], or mass spectrometry [Oeth et al., 2009]). One way to do this is to use only the genotype posterior distributions for each individual, which are often provided by different genotyping software that analyze data from different genotyping technologies [Voorrips et al., 2011, Serang et al., 2012, Gerard et al., 2018, Gerard and Ferrão, 2019, e.g.]. We will thus assume the user provides the posterior means and variances for the genotypes for each individual at two loci, which can be easily obtained from the full posterior distributions for each individual. If genotype posteriors are not provided, genotype likelihoods may be normalized to posterior probabilities (assuming a uniform prior) and used in what follows. The effect of the prior should be negligible for large sample sizes.

To define our estimators of LD, let *X*_*iA*_ and *X*_*iB*_ be the posterior means at loci A and B for individual *i* ∈ {1, …, *n*}. Let *Y*_*iA*_ and *Y*_*iB*_ be the posterior variances at loci A and B for individual *i*. Our estimators are based entirely on the following sample moments of these posterior moments, which may be calculated in linear time in the sample size, *n*.

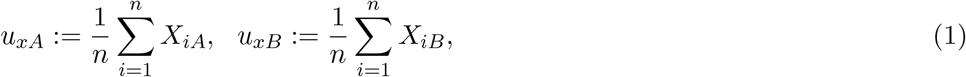

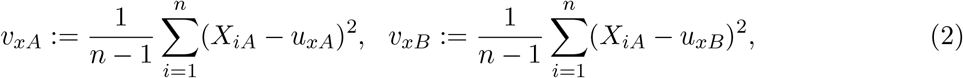

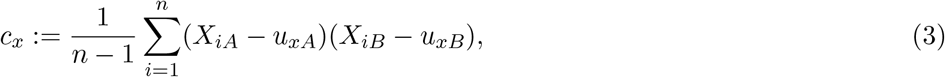

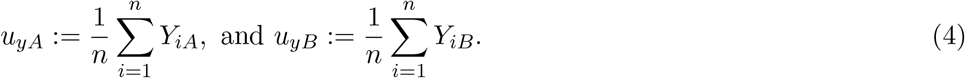

For a *K*-ploid species, our LD estimators, which we derive in Section S1, are as follows. The estimated LD coefficient is

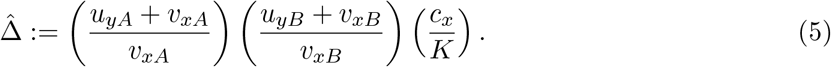

The estimated Pearson correlation is

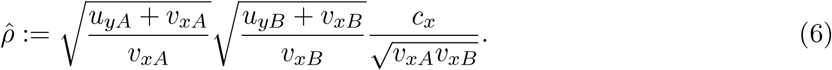

Note that 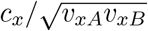 is the sample Pearson correlation between posterior mean genotypes. The estimated standardized LD coefficient is

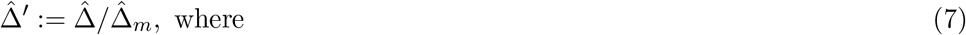

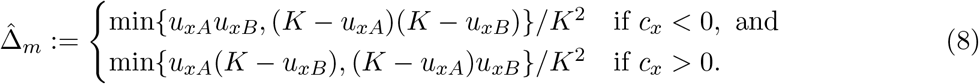

Equations (5)–(7) take the naive estimators most researchers use in practice (the sample covariance/correlation of posterior means) and inflate these by a multiplicative effect. Such multiplicative effects are sometimes called “reliability ratios” in the measurement error models literature [Fuller, 2009]. Due to sampling variability, this inflation could result in estimates that lie beyond the theoretical bounds of the parameters being estimated. In such cases, we apply the following truncations.

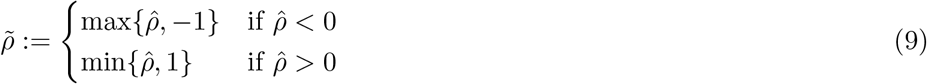

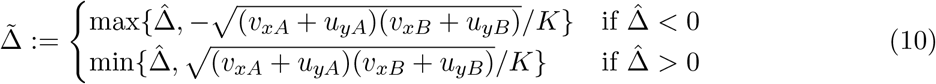

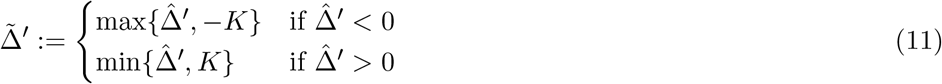

Standard errors are important for hypothesis testing [Brown, 1975], read-depth suggestions [Maruki and Lynch, 2014], and shrinkage [Dey and Stephens, 2018]. Because estimators (5)–(7) are functions of sample moments, deriving their standard errors can be accomplished by appealing to the central limit theorem, followed by an application of the delta method (Section S2).

Additional considerations for improving our estimates of the reliability ratios, such as using hierarchical shrinkage [Stephens, 2016], are considered in Section S3.

All methods are implemented in the `ldsep` package on the Comprehensive R Archive Network https://cran.r-project.org/package=ldsep.

## 3 Results

### 3.1 Simulations

We compared our moment-based estimators (5)–(7) to those of the MLE of Gerard [2021] as well as the naive estimator that calculates the sample covariance and sample correlation between posterior mean genotypes at two loci. Each replication, we generated genotypes for *n* ∈ {10, 100, 1000} individuals with ploidy *K* ∈ {2, 4, 6, 8} under Hardy-Weinberg equilibrium at two loci with major allele frequencies (*p*_*A*_, *p*_*B*_) ∈ {(0.5, 0.5), (0.5, 0.75), (0.9, 0.9)} and Pearson correlation *ρ* ∈ {0, 0.5, 0.9}. We then used `updog`’s `rflexdog()` function [Gerard et al., 2018, Gerard and Ferrão, 2019] to generate read-counts at read-depths of either 10 or 100, a sequencing error rate of 0.01, an overdispersion value of 0.01, and no allele bias. `Updog` was then used to generate genotype likelihoods and genotype posterior distributions for each individual at each SNP. These were then fed into `ldsep` to obtain the MLE, our new moment-based estimator, and the naive estimator. Simulations were replicated 200 times for each unique combination of simulation parameters.

The accuracy of estimating *ρ*^2^ when *p*_*A*_ = *p*_*B*_ = 0.5 at a read-depth of 10 is presented in Figure 1. The results for other scenarios are similar and may be found on Zenodo (https://doi.org/10.5281/zenodo.4543473). We see that the moment-based estimator and the MLE perform comparably, even for small read-depth and sample size. The naive estimator has a strong attenuation bias toward zero. This bias is particularly prominent for higher ploidy levels. For example, for an octoploid species where the true *ρ*^2^ is 0.81, the naive estimator appears to converge to a *ρ*^2^ estimate of around 0.25. This bias does not disappear with increasing sample size. Estimated standard errors are reasonably well-behaved, except for 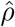 and 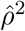 when the sample size is small and the LD is large (Figure 2).

**Figure 1:**
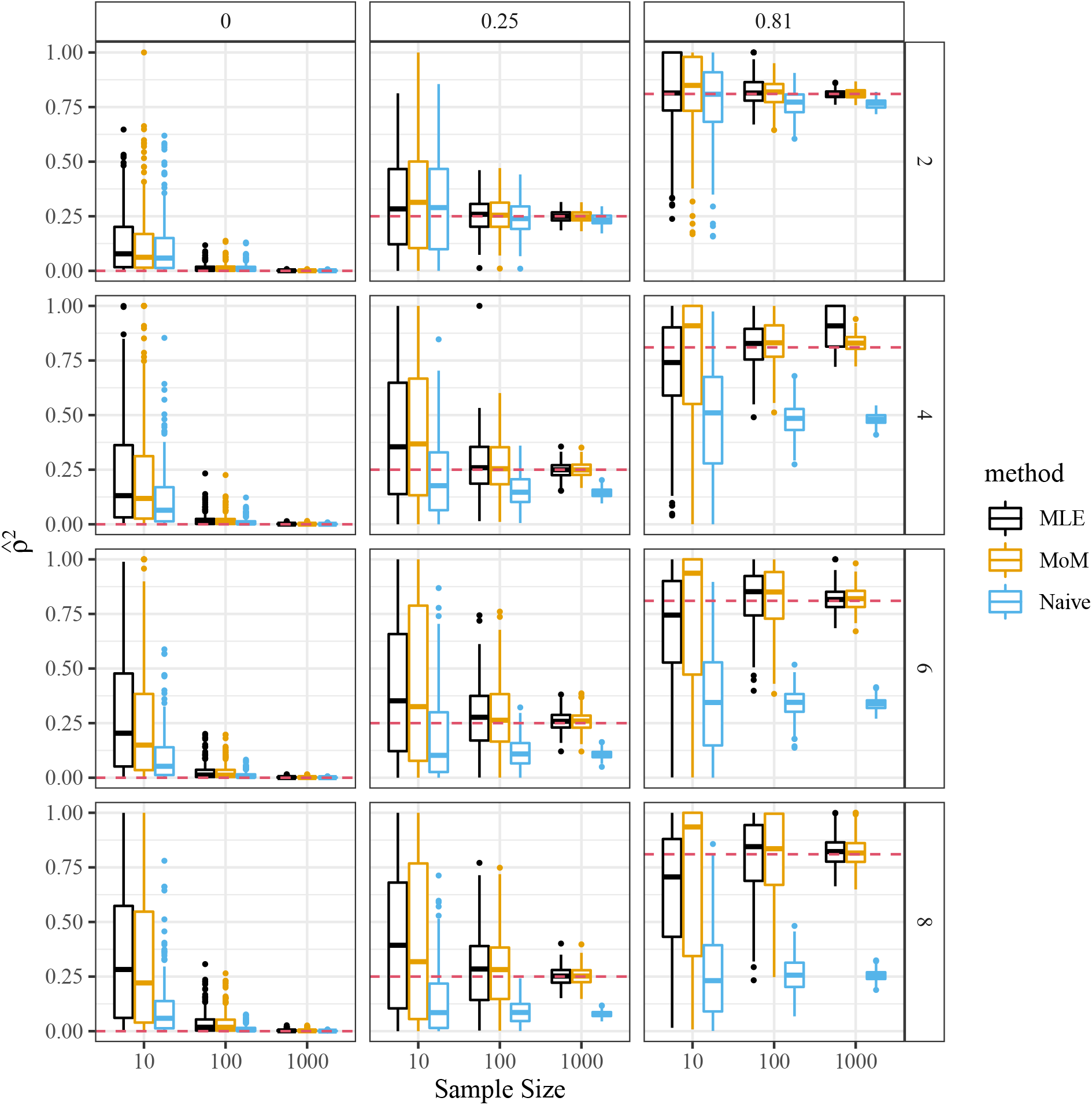
Estimate of *ρ*^2^ (*y*-axis) for the maximum likelihood estimator [Gerard, 2021] (MLE), our new moment-based estimator (6) (MoM), and the naive squared sample correlation coefficient between posterior mean genotypes (Naive). The *x*-axis indexes the sample size, the row-facets index the ploidy, and the column-facets index the true *ρ*^2^, which is also presented by the horizontal dashed red line. These simulations were performed using a read-depth of 10, and major allele frequencies of 0.5 at each locus. The naive estimator presents a strong attenuation bias toward 0, particularly for higher ploidy regimes.

**Figure 2:**
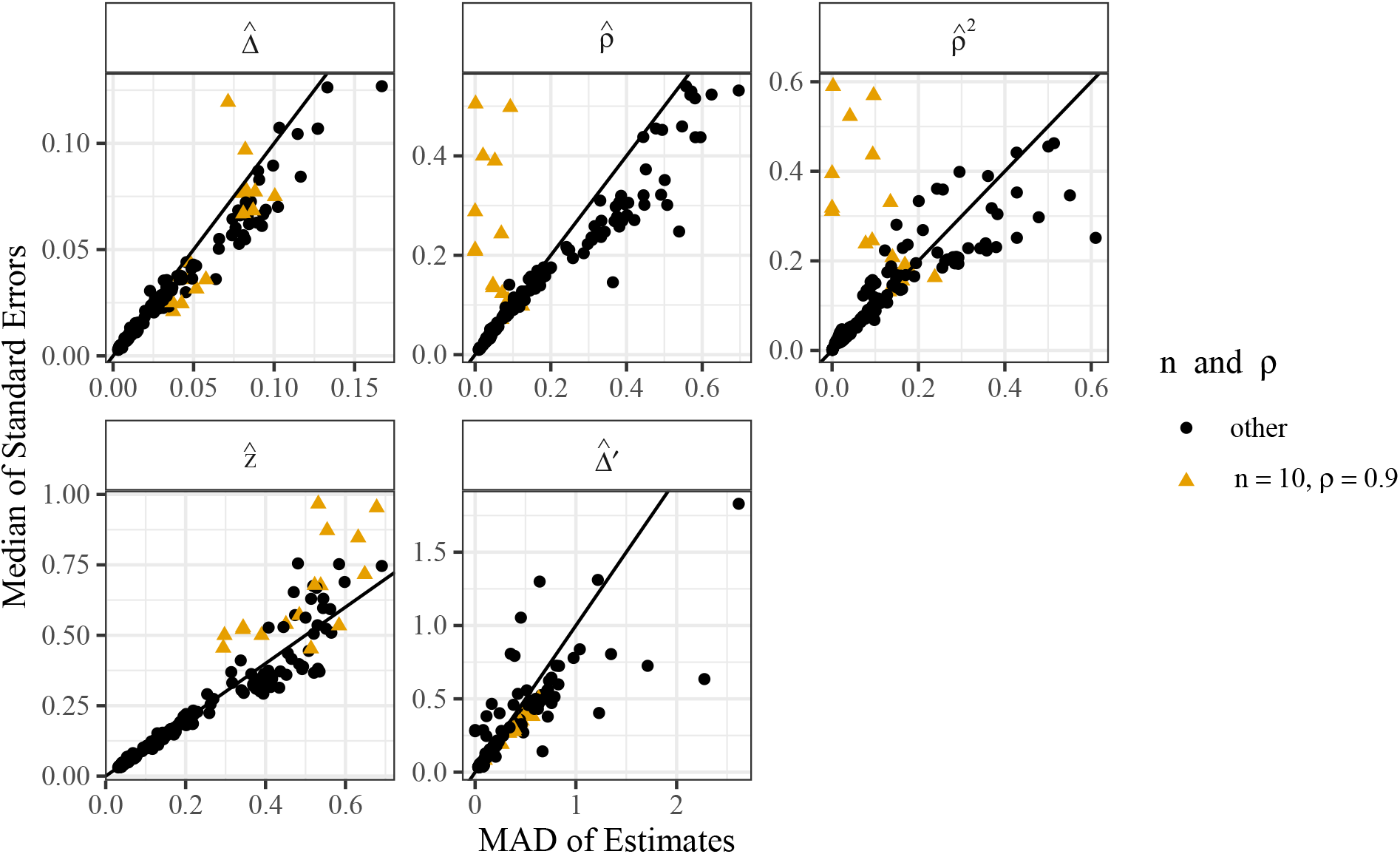
Median of estimated standard errors (*y*-axis) versus median absolute deviations (*x*-axis) of each of the moment-based LD estimators (facets). The line is the *y* = *x* line, and points above this line indicate that the estimated standard errors are typically larger than the true standard errors. Estimated standard error are reasonably unbiased except for 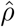 and 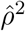 in scenarios with small sample sizes (*n* = 10) and a large levels of LD (*ρ* = 0.9) (color and shape).

### 3.2 LD estimates for *Solanum tuberosum*

We evaluated our methods on the autotetraploid potato (*Solanum tuberosum*, 2*n* = 4*x* = 48) genotyping-by-sequencing data from Uitdewilligen et al. [2013]. We used `updog` [Gerard et al., 2018, Gerard and Ferrão, 2019] to obtain the posterior moments for each individual’s genotype at each SNP on a single super scaffold (PGSC0003DMB000000192). To remove monoallelic SNPs, we filtered out SNPs with allele frequencies either greater than 0.95 or less than 0.05, and filtered out SNPs with a variance of posterior means less than 0.05. This resulted in 2108 SNPs. We then estimated the squared correlation between each SNP using either the naive approach of calculating the sample Pearson correlation between posterior means, or using our new moment-based approach (6).

Our estimators are scalable. On a 1.9 GHz quad-core PC running Linux with 32 GB of memory, it took a total of 1.9 seconds to estimate *all* pairwise correlations using our new moment-based approach, which is a small increase over the 0.7 seconds it took to estimate all pairwise correlations using the naive approach. In Gerard [2021], we found that the MLE approach took about 0.1 seconds for *each* pair of SNPs for a tetraploid individual. Extrapolating this to 2108 SNPs would indicate that the MLE approach would take about 2.5 days of computation time to calculate all pairwise LD estimates on this dataset 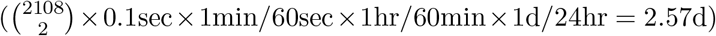.

The histogram of estimated reliability ratios are presented in Figure 3. We see there that the reliability ratios of most SNPs only increase their correlation estimates by less than 10%. But a not insignificant portion have reliability ratios that increase the correlation estimates by more than 10%. To evaluate the LD estimates of high reliability ratio SNPs, we calculated the MLEs for *ρ*^2^ between the twenty SNPs with the largest reliability ratios. A pairs plot for *ρ*^2^ estimates between the three approaches is presented in Figure 4. We see there that the MLE and new moment-based approach result in very similar *ρ*^2^ estimates, while the naive approach using posterior means results in much smaller *ρ*^2^ estimates.

**Figure 3:**
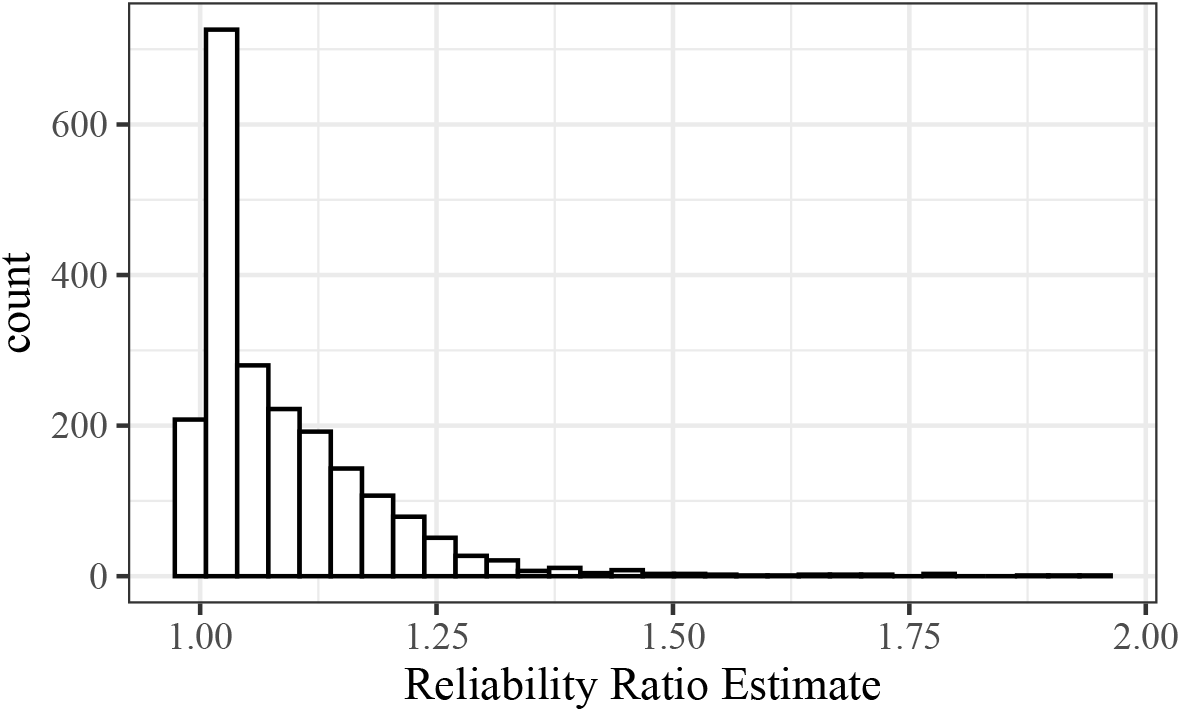
Histogram of estimated reliability ratios (S69) using the data from Uitdewilligen et al. [2013].

**Figure 4:**
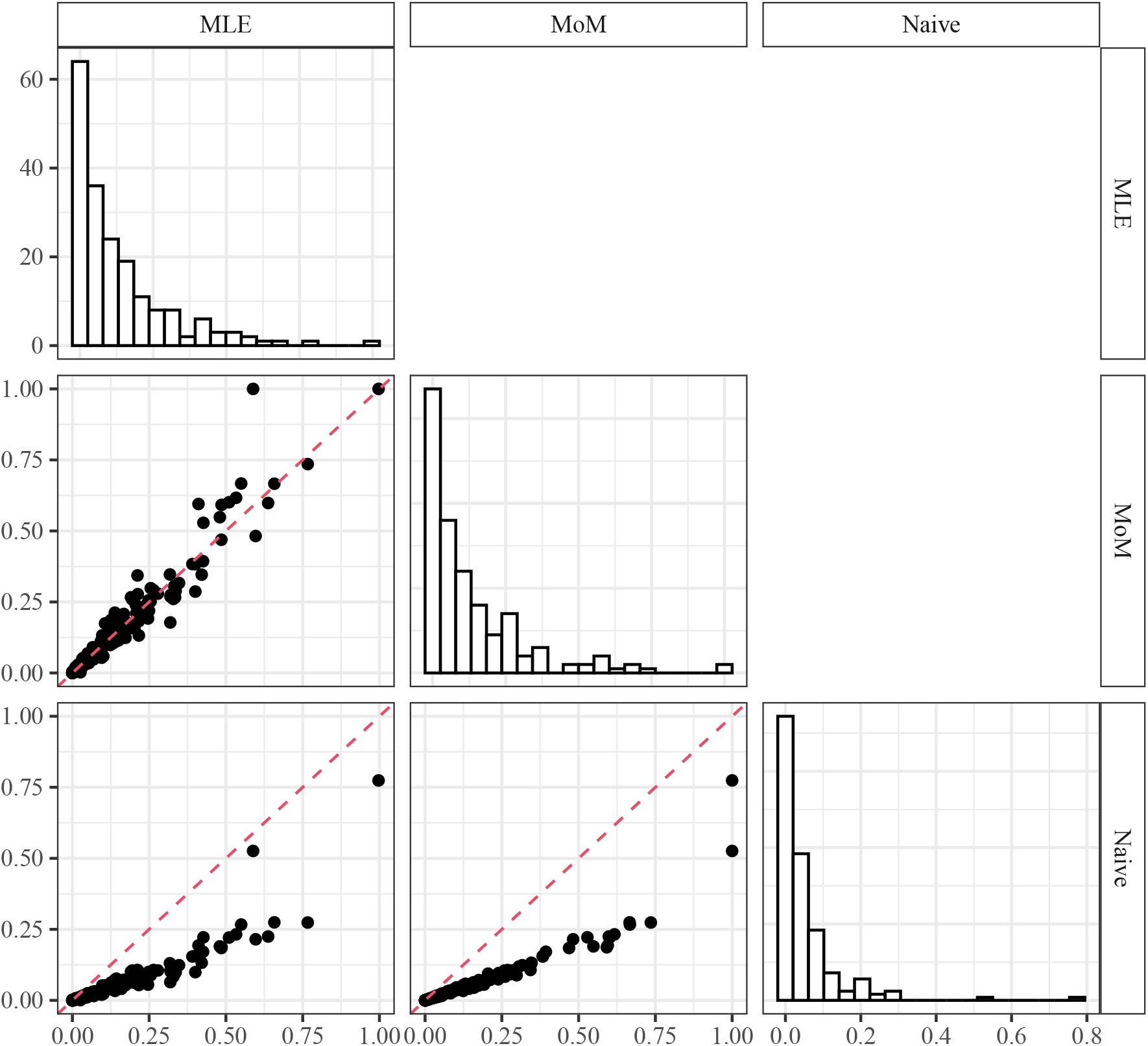
Pairs plot for *ρ*^2^ estimates between the twenty SNPs from Uitdewilligen et al. [2013] with the largest estimated reliability ratios when using either maximum likelihood estimation (MLE) [Gerard, 2021], our new moment-based approach (6) (MoM), or the naive approach using just posterior means (Naive). The dashed line is the *y* = *x* line. The MLE and the moment-based approach result in much more similar LD estimates.

## 4 Discussion

It has been known since at least the time of Spearman that the sample correlation coefficient (or, similarly, the ordinary least squares estimator in simple linear regression) is attenuated in the presence of uncertain variables [Spearman, 1904]. Methods to adjust for this bias include assuming prior knowledge on the measurement variances or the ratio of measurement variances (resulting from, for example, repeated measurements on the same individuals) [Koopmans, 1937, Degracie and Fuller, 1972], using instrumental variables [Carter and Fuller, 1980], and using distributional assumptions [Pal, 1980]. See Fuller [2009] for a detailed introduction to this vast field. In order to accommodate different data types [Fan et al., 2003, Baird et al., 2008, Oeth et al., 2009, Elshire et al., 2011] and different genotyping programs [Voorrips et al., 2011, Serang et al., 2012, Gerard et al., 2018, Gerard and Ferrão, 2019], and therefore increase the generality of our methods, we limited ourselves to using just posterior genotype probabilities to calculate LD. This excluded using these previous approaches. Our solution, then, was to use sample moments of marginal posterior moments which, to our knowledge, has never been proposed before.

It is natural to ask if our methods could be used to account for uncertain genotypes in genomewide association studies. However, the moment-based techniques we used in this manuscript, when applied to simple linear regression with an additive effects model (where the SNP effect is proportional to the dosage), result in the standard ordinary least squares estimates when using the posterior mean as a covariate (Section S4). This supports using the posterior mean as a covariate in simple linear regression with an additive effects model. This is not to say, however, that using the posterior mean is also appropriate for more complicated models of gene action [Rosyara et al., 2016], or for non-linear models.

## Acknowledgments

Most analyses were performed using the R statistical language [R Core Team, 2020].

## Data availability

All methods discussed in this manuscript are implemented in the `ldsep` package, available on the Comprehensive R Archive Network (https://cran.r-project.org/package=ldsep) under a GPL-3 license. Scripts to reproduce the results of this research are available on Zenodo (https://doi.org/10.5281/zenodo.4543473). All datasets used in this manuscript are publicly available [Uitdewilligen et al., 2013] and may be downloaded from:

- https://doi.org/10.1371/journal.pone.0062355.s004
- https://doi.org/10.1371/journal.pone.0062355.s007
- https://doi.org/10.1371/journal.pone.0062355.s009
- https://doi.org/10.1371/journal.pone.0062355.s010

## Supplementary Material

### S1 Derivation of LD estimators

In this section, we derive estimators (5)–(7). We do this by assuming a normal model on the data and the genotypes. This is obviously not appropriate when using genotypes and sequencing data, but our simulations in Section 3.1 were also accomplished using sequencing data and resulted in very good performance.

Let ***G***_*i*_ = (*G*_*iA*_, *G*_*iB*_)^T^ be the genotype for individual *i* at loci A and B. Let ***Z***_*i*_ = (*Z*_*iA*_, *Z*_*iB*_)^T^ be the data for individual *i* at loci A and B. Then we let

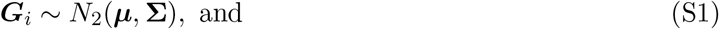

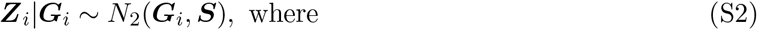

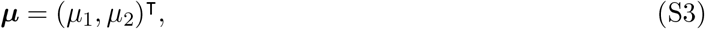

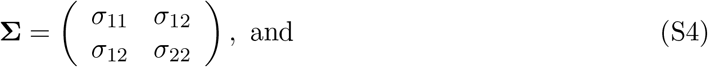

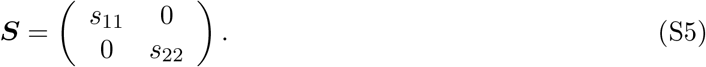

To interpret these terms, *µ*_1_*/K* and *µ*_2_*/K* are the allele frequencies at each locus, *σ*_11_ and *σ*_22_ are the variances of the genotypes at each locus, *s*_11_ and *s*_22_ are the variances of the genotyping errors at each locus, and *σ*_12_ is the covariance between genotypes. By elementary methods, we have the well-known result that, marginally,

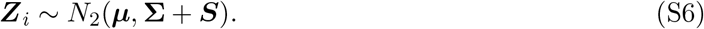

We assume the user has provided posterior moments on the genotypes

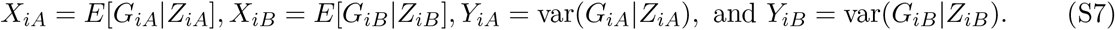

These posterior moments are marginal in that they only condition on either *Z*_*iA*_ or *Z*_*iB*_, but not both. Thus, we assume they are well-approximated by the model

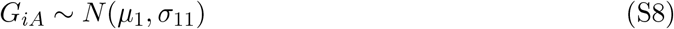

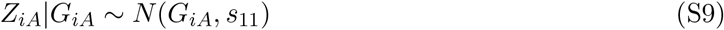

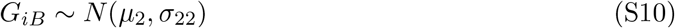

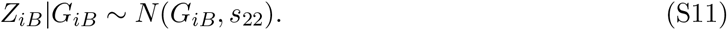

By standard methods, this results in

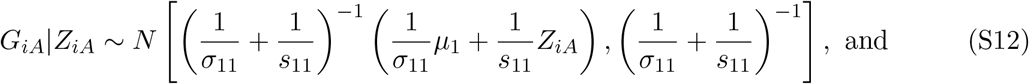

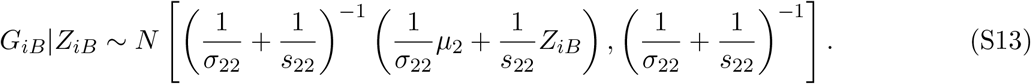

Treating only ***Z***_*i*_ as random from distribution (S6), we have

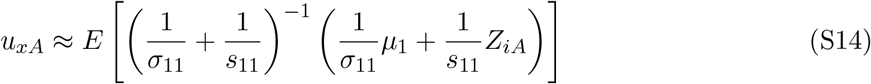

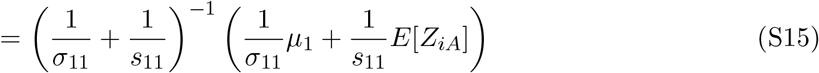

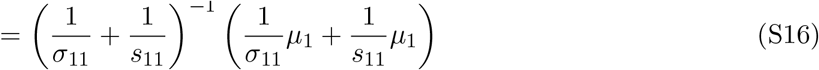

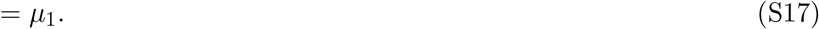

Similarly,

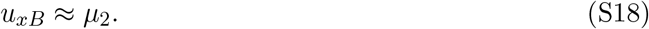

Furthermore,

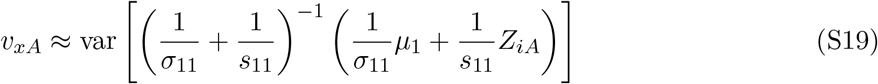

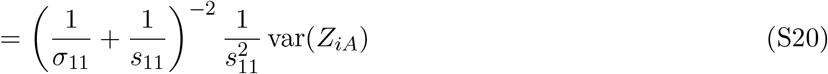

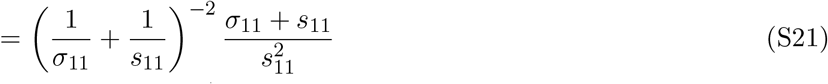

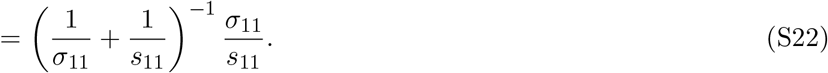

Similarly,

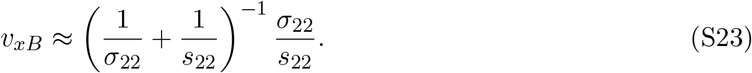

Now, using the posterior variances, we have

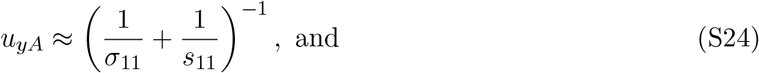

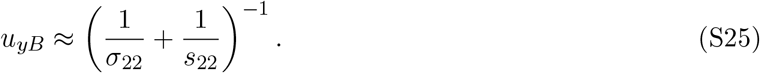

Finally, the expectation of the sample covariance of posterior means is

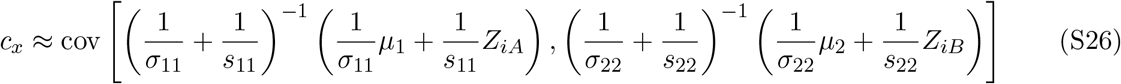

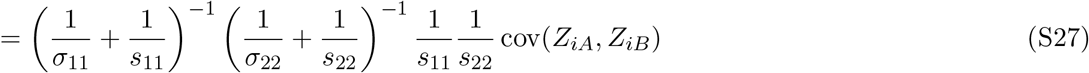

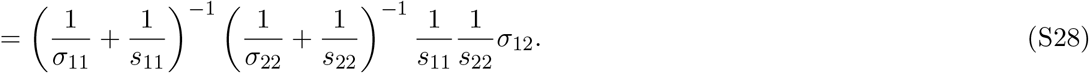

Using a method-of-moments approach, we now have a system of five equations and five unknowns:

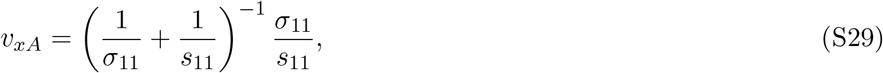

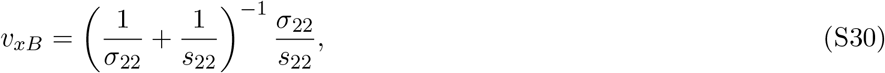

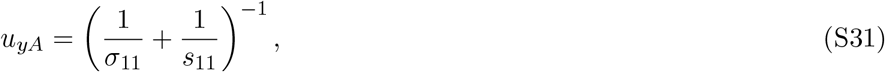

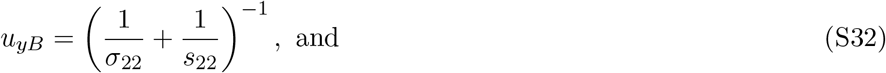

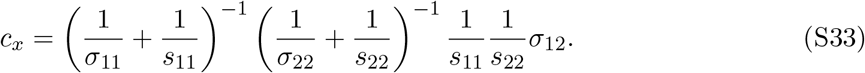

Solving for *s*_11_, *s*_22_, *σ*_11_, *σ*_22_, and *σ*_12_, we obtain:

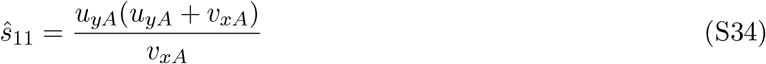

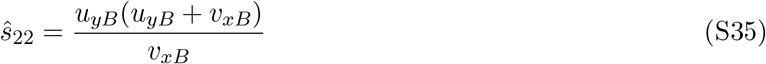

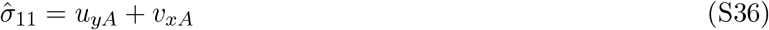

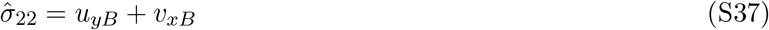

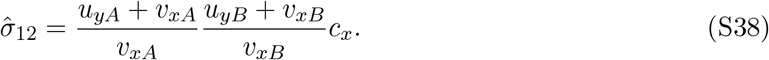

Using (S14)–(S18), we also have

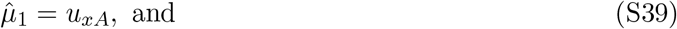

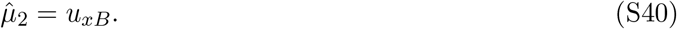

The LD coefficient estimates (5)–(7) can be obtained by substituting in parameter estimates in the following equations [Gerard, 2021]

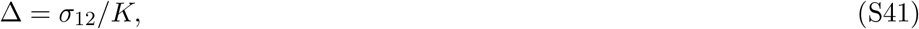

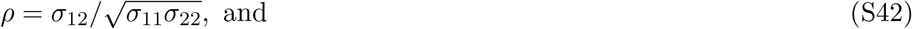

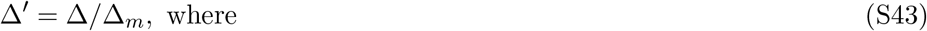

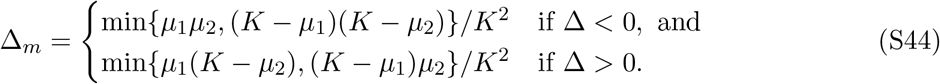

### S2 Derivation of standard errors

Let

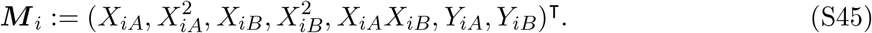

Then, by the central limit theorem, we have for

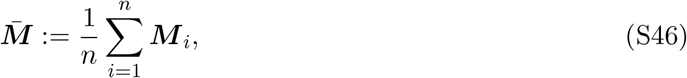

that 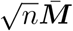 is asymptotically multivariate normal with some limiting covariance, say, **Ω**. Finite variances are guaranteed by the finite support of the genotypes. We can estimate **Ω** with the sample covariance matrix

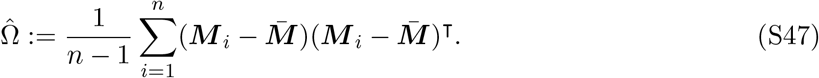

Estimators (5)–(7) are approximately functions of 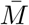. Namely

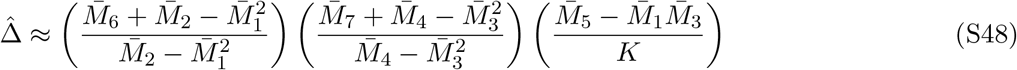

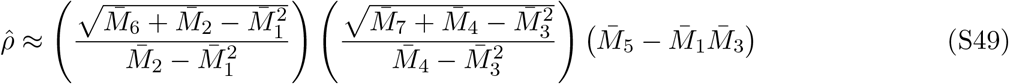

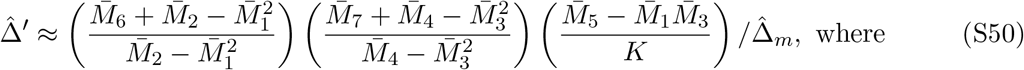

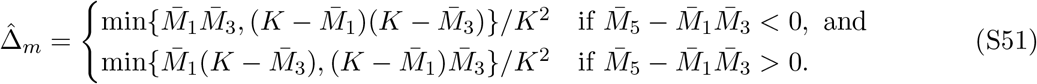

These are smooth functions of 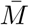 (except on a space of Lebesgue measure zero), and so admit the following gradients, calculated in Mathematica [Wolfram Research, Inc., 2020]:

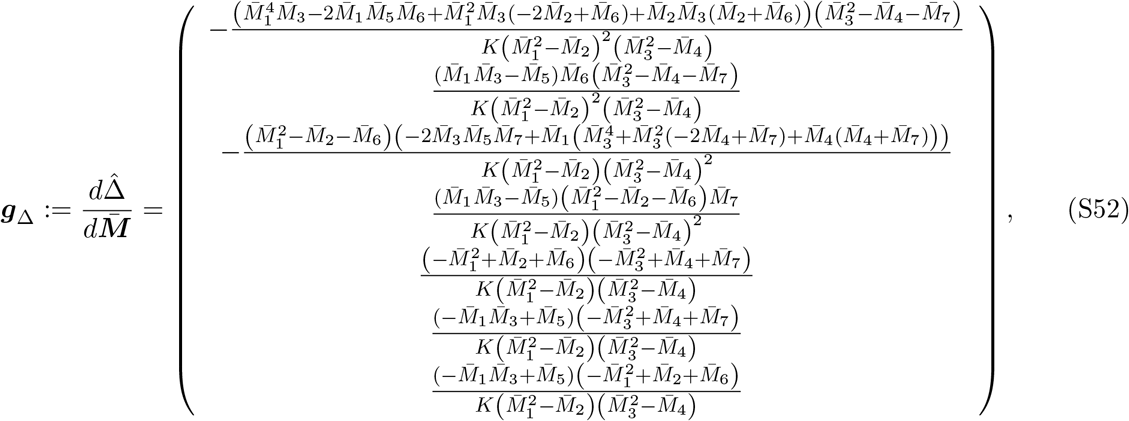

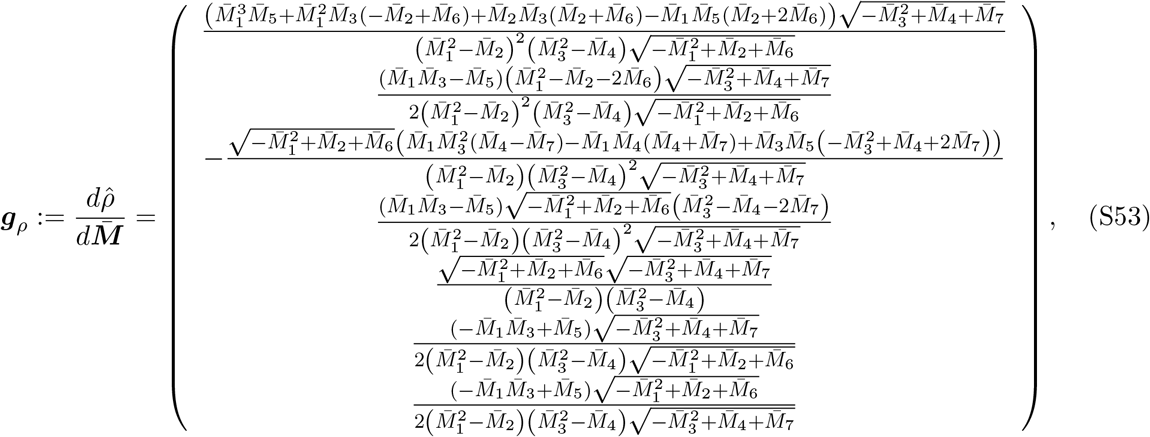

and

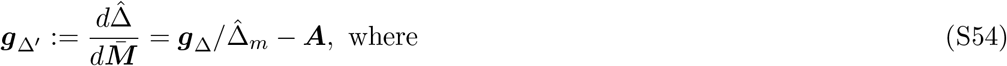

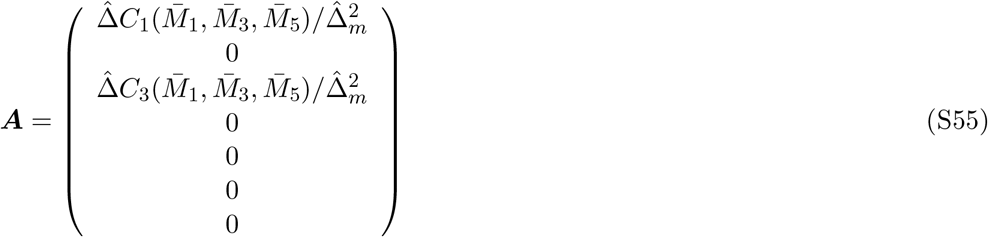

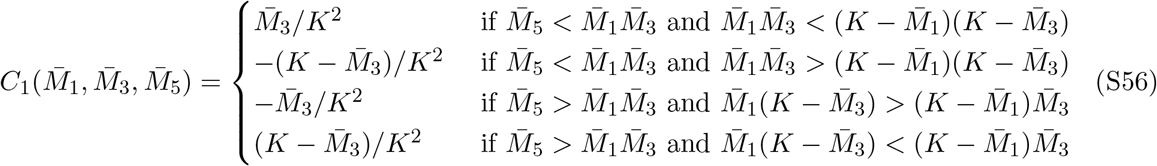

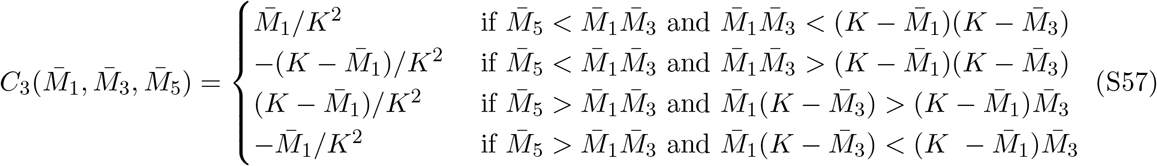

Though these gradients are rather complicated, they are not computationally intensive and may be calculated in constant time in the sample size.

The asymptotic variances of 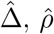, and 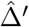 are

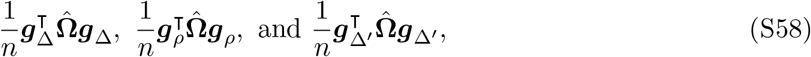

respectively.

To accommodate missing data, we use only pairwise complete observations for the sample covariance matrix (S47). This ensures that 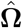 is positive definite and, thus, the resulting standard errors are non-negative. However, we use all non-missing observations for 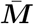 That is, let Θ_*A*_, Θ_*B*_ ⊆ {1, 2, …, *n*} be the index sets of non-missing values at loci A and B, respectively. Then

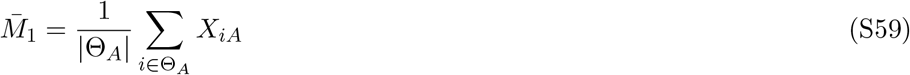

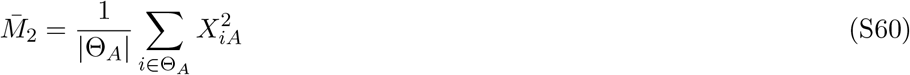

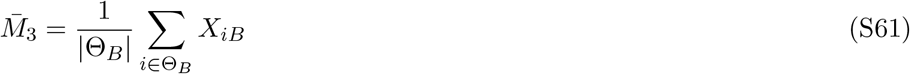

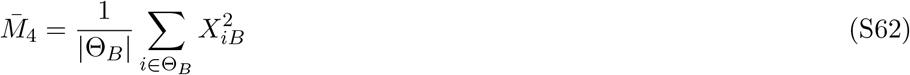

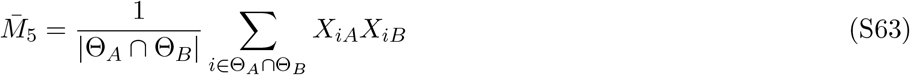

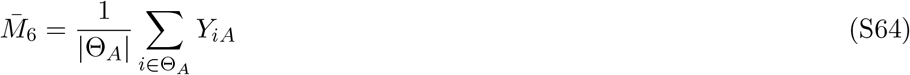

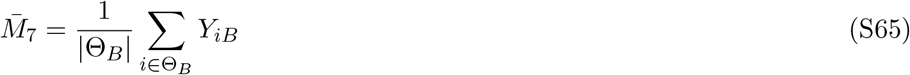

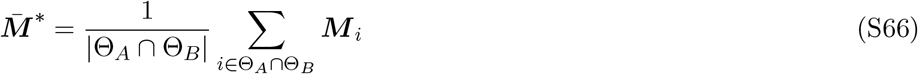

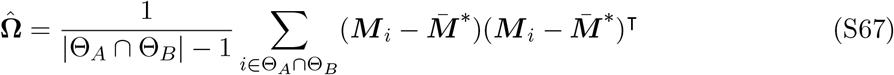

The asymptotic variances of 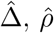, and 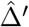 are then

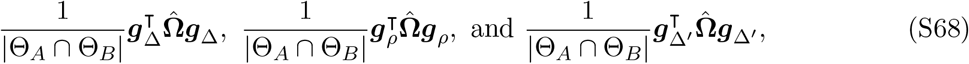

respectively.

### S3 Adjusting the reliability ratios

#### S3.1 Adaptive shrinkage on the reliability ratios

Each SNP has an estimated reliability ratio,

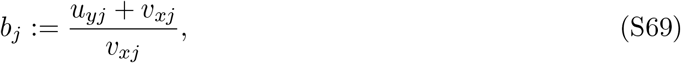

which corresponds to the multiplicative adjustment to all LD estimates that include that SNP (see (5)). These reliability ratios might have high variance due to (i) lower sequencing depth or (ii) containing fewer individuals with non-missing data. Thus, some reliability ratios may be noisy. Hierarchical shrinkage is a statistical technique that allows high-variance observations to borrow strength from low-variance observations and thus improve estimation performance. Adaptive shrinkage (*ash*) [Stephens, 2016] is a recently proposed general-purpose hierarchical shrinkage technique that we can use to model the distribution of reliability ratios flexibly, only constraining them to be unimodal. In this section, we will use *ash* to improve our reliability ratio estimates.

We will now describe the procedure for applying *ash* to shrink the reliability ratios. Our strategy will be to derive the standard errors for the log of the reliability ratios (S69) and apply *ash* on the log-scale using these standard errors. To begin, let *X*_*ij*_ be the posterior mean for individual *i* at SNP *j*. Let *Y*_*ij*_ be the posterior variance for individual *i* at SNP *j*. Finally, let

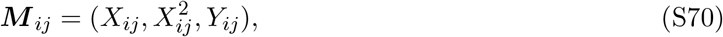

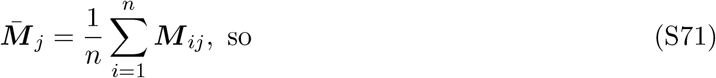

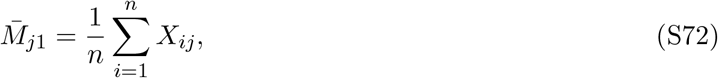

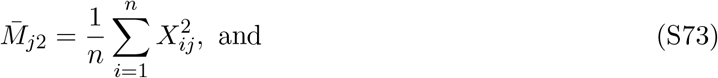

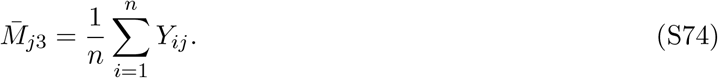

Then the log of the reliability ratio for SNP *j* is

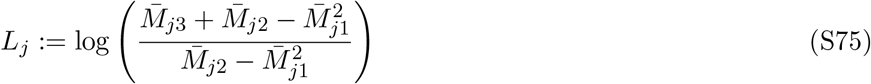

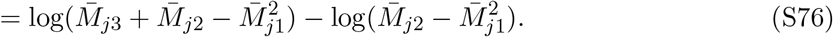

Let the sample covariance be

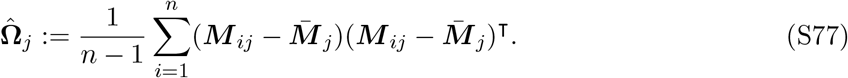

Then we have by the central limit theorem that 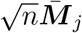 is asymptotically multivariate normal, and we can use 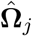 as the estimate of the covariance matrix. The gradients for (S75) are

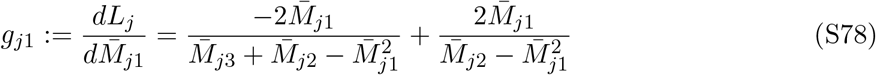

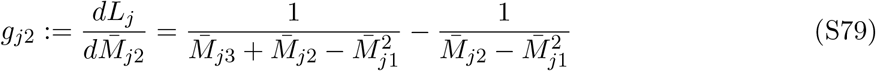

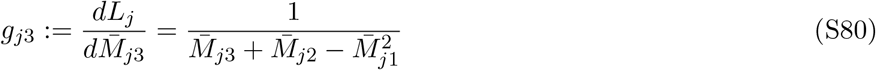

Then, with *g*_*j*_:= (*g*_*j*1_, *g*_*j*2_, *g*_*j*3_)^T^, the variance for *L*_*j*_ is

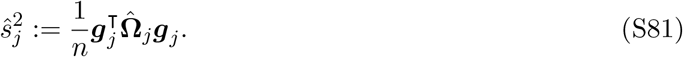

We apply *ash* to (*L*_1_, ŝ_1_), …, (*L*_*m*_, ŝ_*m*_) to obtain shrunken log reliability ratios 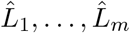. Because *ash*’s grid-based scheme for estimating the mode is not the most computationally efficient, we used the half-sample mode estimator of Robertson and Cryer [1974] prior to running *ash*.

This procedure seems to result in improved performance for SNPs with unusually variable reli-ability ratios (Figure S1).

#### S3.2 Thresholding the reliability ratios

If a researcher accidentally provides a monoallelic SNP, its reliability ratio could explode due to having a denominator close to zero in (S69). For example, the right panel of Figure S2 contains a monoallelic SNP (PotVar0080327) whose reliability ratio estimate (S69) is 100.92. This can provide unstable estimates of LD as some SNPs will, due to sampling variability, have correlations with these monoallelic SNPs on the order of 0.01. For example, the sample correlation between posterior means of PotVar0080327 and PotVar0078678 (left facet of Figure S2) −0.0098. But due to the extreme reliability ratio of PotVar0080327, the genotype-error adjusted correlation estimate is −1. This is, of course, unsettling. So by default, our software will take all reliability ratio estimates (S69) above a user-provided value (default of 10) and assign these to have reliability ratios of the median reliability ratio in the dataset.

### S4 Genome-wide Association Studies

In this section, we demonstrate that the techniques used in Section S1, when applied to simple linear regression with an additive effects model [Rosyara et al., 2016], result in the standard ordinary least squares estimate when using the posterior mean as a covariate. This indicates that for genome-wide association studies, using the posterior mean is appropriate in a linear regression context when using an additive model for gene action.

Let *G*_*i*_ be the genotype for individual *i* at a locus. Let *Z*_*i*_ be the data that lead to the genotyping for individual *i* at the same locus. Let *W*_*i*_ be some quantitative trait of interest for individual *i*. Then we let

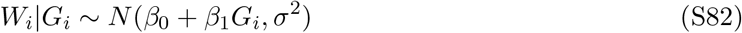

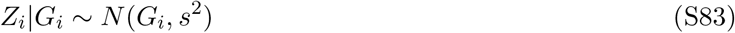

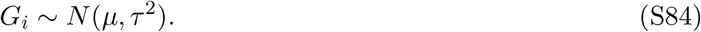

We suppose the user is only provided the posterior means and variances of each *G*_*i*_|*Z*_*i*_. Let *X*_*i*_ =*E*[*G*_*i*_|*Z*_*i*_] and *Y*_*i*_ = var(*G*_*i*_|*Z*_*i*_). From elementary methods, we have

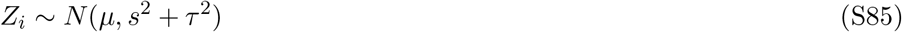

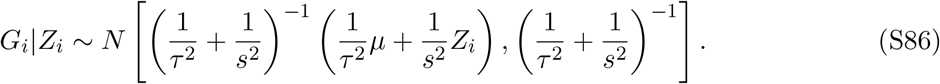

Let

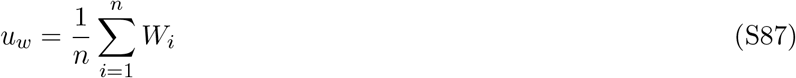

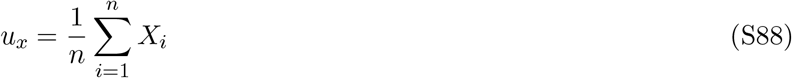

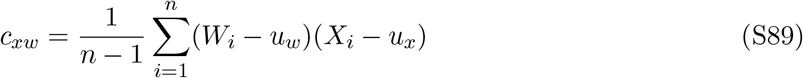

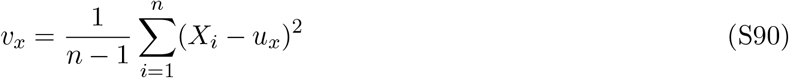

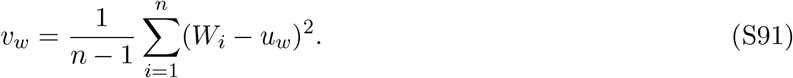

We have that

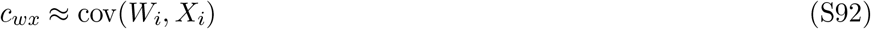

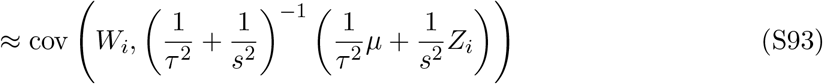

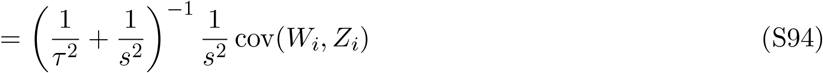

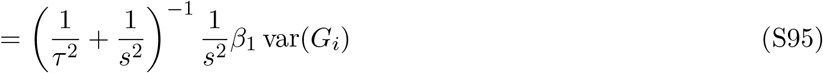

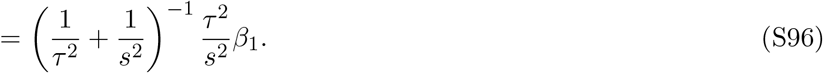

We also have from (S19)–(S22) that

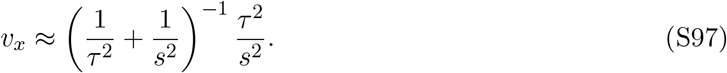

Using method of moments with equations (S96) and (S97), we have the following estimator for *β*_1_

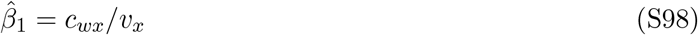

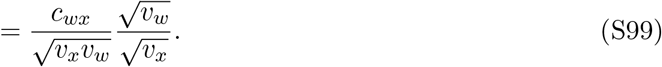

Equation (S99) is the sample correlation between the *W*_*i*_’s and the 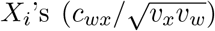 multiplied by the ratio of the sample standard deviations of the *W*_*i*_’s and the 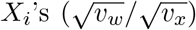. This is the well-known formula for the ordinary least squares estimate of *β*_1_ from a regression of *W*_*i*_ on *X*_*i*_.

### S5 Supplementary figures

**Figure S1:**
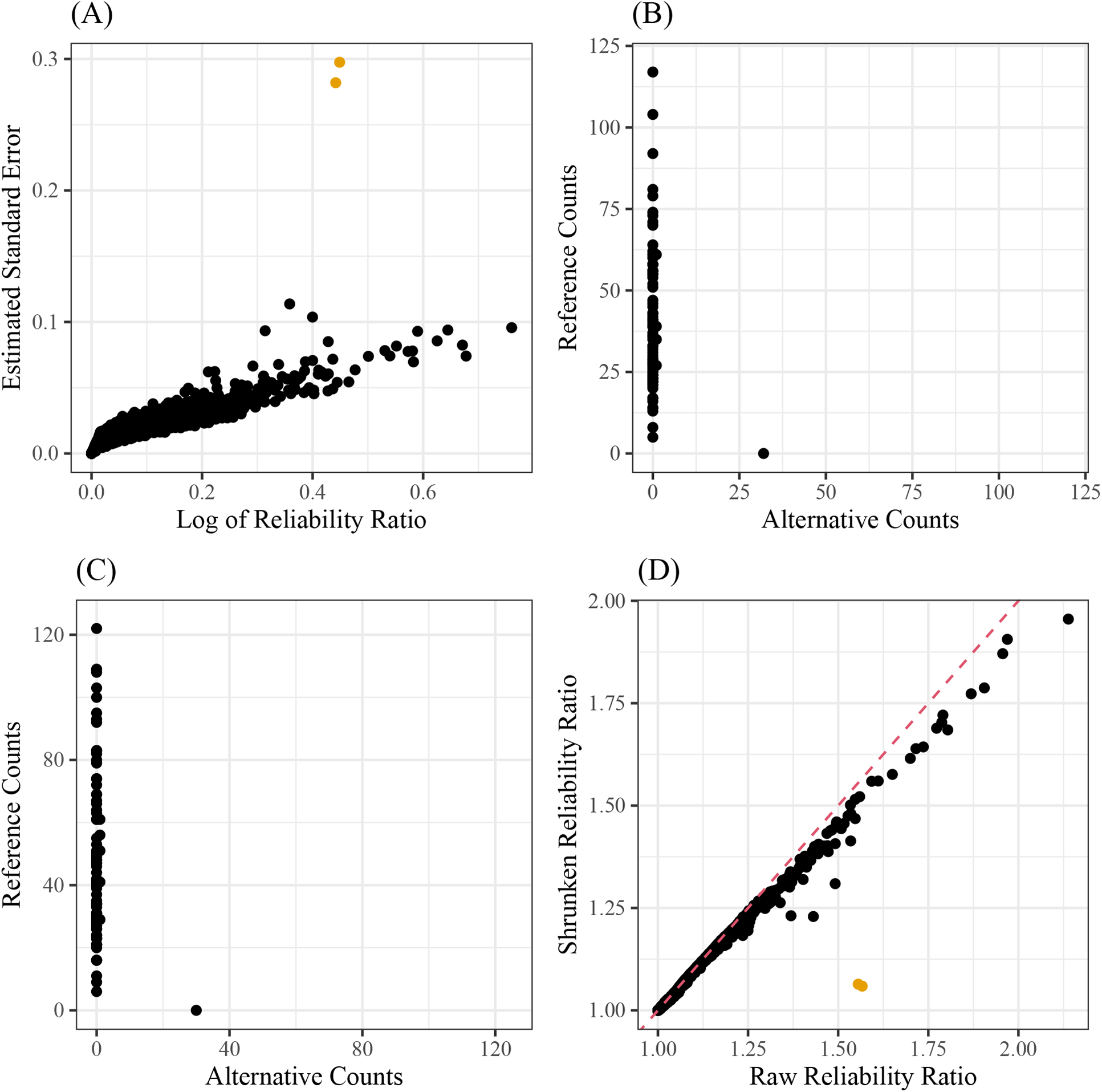
**(A)** The log of the reliability ratios (*x*-axis) versus their estimated standard errors (*y*-axis). The two highlighted points do not seem to fit the trend. When we plot the read-counts for these highlighted points (**(B)** and **(C)**), we notice that these two SNPs are almost monoallelic, providing doubts on their unusually large reliability ratios. We plot the shrunken reliability ratios (*y*-axis) against their original values (*x*-axis) in **(D)**, noting that the problem SNPs (color) have their reliability ratios highly adjusted.

**Figure S2:**
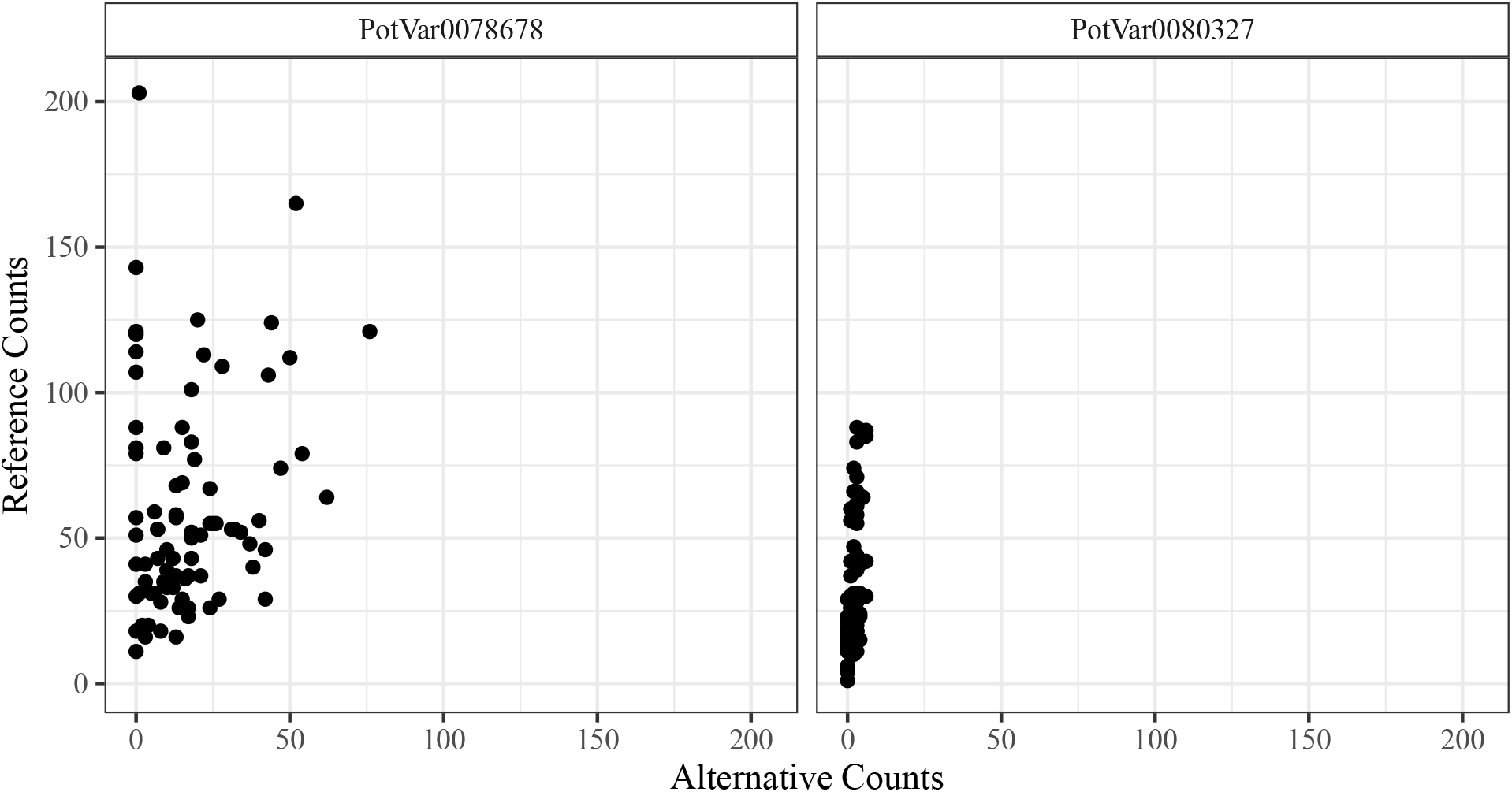
Plots of read-counts of two SNPs (facets) from Uitdewilligen et al. [2013]. Alternative counts lie on the *x*-axis and reference counts lie on the *y*-axis. The right SNP is monoallelic and because of this the estimated correlation between the two SNPs using raw reliability ratios is −1, even though the sample correlation between posterior means is only −0.0098.

